# Incomplete protection against dengue virus type 2 re-infection in Peru

**DOI:** 10.1101/023747

**Authors:** Brett M. Forshey, Robert C. Reiner, Sandra Olkowski, Amy C. Morrison, Angelica Espinoza, Kanya C. Long, Stalin Vilcarromero, Wilma Casanova, Helen J. Wearing, Eric S. Halsey, Tadeusz J. Kochel, Thomas W. Scott, Steven T. Stoddard

## Abstract

**Background.** Nearly half of the world’s population is at risk for dengue, yet no licensed vaccine or anti-viral drug is currently available. Dengue is caused by any of four dengue virus serotypes (DENV-1 through DENV-4), and infection by a DENV serotype is assumed to provide life-long protection against re-infection by that serotype. We investigated the validity of this fundamental assumption during a large dengue epidemic caused by DENV-2 in Iquitos, Peru, in 2010-2011, 15 years after the first outbreak of DENV-2 in the region.

**Methodology/Principal Findings.** We estimated the age-dependent prevalence of serotype-specific DENV antibodies from longitudinal cohort studies conducted between 1993 and 2010. During the 2010-2011 epidemic, active dengue cases were identified through active community- and clinic-based febrile surveillance studies, and acute inapparent DENV infections were identified through contact tracing studies. Based on the age-specific prevalence of DENV-2 neutralizing antibodies, the age distribution of DENV-2 cases was markedly older than expected. Homologous protection was estimated at 35.1% (95% confidence interval: 0% -- 65.2%). At the individual level, pre-existing DENV-2 antibodies were associated with an incomplete reduction in the frequency of symptoms. Among dengue cases, 43% (26/66) exhibited elevated DENV-2 neutralizing antibody titers for years prior to infection, compared with 76% (13/17) of inapparent infections (age-adjusted odds ratio: 4.2; 95% confidence interval: 1.1 – 17.7).

**Conclusions/Significance.** Our data indicate that protection from homologous DENV re-infection may be incomplete in some circumstances, which provides context for the limited vaccine efficacy against DENV-2 in recent trials. Further studies are warranted to confirm this phenomenon and to evaluate the potential role of incomplete homologous protection in DENV transmission dynamics.

**Author Summary:** Homotypic immunity against DENV infection has been assumed to be complete and lifelong, and to our knowledge, instances of homologous DENV re-infection have not been rigorously documented. However, few long-term studies have been conducted in such a way that homologous re-infection could be observed, if it did in fact occur. Our study provides evidence that homologous re-infection may occur in certain circumstances. We draw from data collected during a 2010-2011 DENV-2 epidemic in northeastern Peru, 15 years after the initial DENV-2 outbreak in the region. This finding has significant implications for our understanding of dengue epidemiology and for dengue vaccine formulation, which may need to consider multiple genotypes of each serotype. Data from other long-term dengue epidemiology studies should be analyzed to determine of homologous re-infection is a more widespread phenomenon.

## Introduction

Dengue is a mosquito-borne viral illness that imposes a tremendous public health burden on tropical and sub-tropical regions. An estimated 390 million infections occur globally each year, and up to 4 billion people are at risk [1]. Dengue is caused by four dengue virus (DENV) serotypes (DENV-1 to DENV-4). Infection with any DENV can lead to a range of disease outcomes, from mild febrile illness to severe, hemorrhagic manifestations and death. Although DENV infections are often inapparent, many dengue cases require hospitalization, which can overwhelm medical infrastructure during epidemics. There are no specific antiviral therapeutics and currently no licensed vaccine.

DENVs display substantial inter-serotypic genetic heterogeneity. Serotypes share less than 70% identity at the nucleotide level and less than 80% identity at the amino acid level [2], similar to the genetic distance between Japanese encephalitis virus and West Nile virus. Infection by one DENV serotype appears to induce relatively short term protection against infection by a heterologous serotype [3,4]. Thereafter, an individual returns to being susceptible to heterologous infection [5,6]. Importantly, an individual’s specific DENV infection history can either enhance or attenuate the severity of disease they experience during subsequent, sequential exposures [7–9].

Although DENVs are genetically and phenotypically diverse within serotypes, infection with one strain of a serotype is thought to induce lifelong protection against infection by all other strains of the homologous serotype [10]. This assumption, however, lacks direct support, in part because of difficulty in obtaining the appropriate epidemiological data and the absence of an adequate animal model for studying DENV pathogenesis. In neutralization assays, serum and monoclonal antibody titers can vary markedly depending on the viral strain used [11], which suggests that intra-serotype variability could be epidemiologically important. If re-infection with a new strain of a homologous DENV serotype can produce symptomatic disease, even if tempered by imperfect neutralization, this would have significant ramifications for the ongoing development of DENV vaccines [12] and our understanding of DENV transmission dynamics.

Between late 2010 and early 2011, Iquitos, Peru, experienced an unprecedented outbreak of severe dengue caused predominantly by an American/Asian genotype of DENV-2 (AA-DENV-2) [13,14]. The number of cases (~25,000) reported by the local health authorities [15] far exceeded any previous epidemic in Iquitos, with greater than 2,500 cases per week reported on several occasions. Many cases required hospitalization, which overwhelmed the local health care system [16]. This epidemic occurred 15 years after DENV-2 (American genotype; Am-DENV-2) was first detected in Iquitos, which caused a large outbreak of febrile illness starting in 1995. In the intervening years between the two outbreaks, few isolates of DENV-2 were detected in Iquitos [17]. The 1995 introduction of Am-DENV-2 caused a large outbreak of febrile illness and led to a high prevalence of DENV-2 antibodies among Iquitos residents [18,19]. Given the high DENV-2 antibody prevalence and the sheer magnitude of the dengue outbreak, we postulated that the pre-existing antibodies failed to protect against reinfection and disease. We tested this hypothesis by analyzing population- and individual-level infection patterns, utilizing data from ongoing clinic-and community-based febrile surveillance and a long-term series of longitudinal studies on the seroprevalence of DENV in Iquitos.

## Methods

Ethical considerations. All human subjects protocols (see [20] for protocol numbers) were in compliance with all applicable US Federal regulations governing the protections of human subjects. All protocols received approval from the institutional review boards of all participating institutions. In addition, protocols were reviewed and approved by the Loreto Regional Health Department, which oversees health-related research in Iquitos. Written consent was provided by adult study participants or by parents or guardians of participating minors.

Prospective longitudinal cohorts. Samples used for estimating serotype-specific antibody prevalence between 1993 and 2010 were collected from five different longitudinal cohort studies [20]. Cohort participation was restricted to participants above the age of 5; four of the cohorts included adult participants above the age of 17. Serum samples were collected at 6 to 12 month intervals and were tested for DENV neutralizing antibodies by the plaque reduction neutralization test (PRNT). In the first study (the source data for years 1993 (n=283), 1994 (n=195), 1995 (n=162), and 1996 (n=333)), yearly serum samples were collected from school-aged children (5 – 22 years) who attended six schools in Iquitos [18]. The second cohort (under the study name “Entomological Correlates of Dengue Control”; the source data for years 1999 (n=1,444) and 2002 (n=2,055), and part of the source data for 2004 (combined n=3,068)) was initiated in 1999, continued through 2005 [19], and included a geographically stratified sample of approximately 2,400 participants (school age children and their family members). After the base cohort was established, follow-up samples were collected at approximately 6-month intervals. The third cohort (study name “Dengue Vector Control System”; part of the source data for 2004 (combined n=3,068)) was initiated in 2004, with 2,415 participants providing a baseline serum sample [19]. Follow-up serum samples were collected in January—February 2005 and October 2005. The fourth cohort (study name “Active Dengue Surveillance and Predictors of Disease Severity in Iquitos, Peru”; the source data for 2006 (n=2,356), and part of the source data for years 2008 (combined n=2,137) and 2010 (combined n=3,131)) was initiated in 2006, and included approximately 2,400 participants at baseline. Samples were collected approximately every six months until mid-2010. The final cohort (“Measuring Entomological Risk for Dengue”; part of the source data for years 2008 (combined n=2,137) and 2010 (combined n=3,131)) was initiated in late 2007 among approximately 2,400 participants [21] and is ongoing. Individuals above the age of five years residing in study neighborhoods were invited to participate. Longitudinal serum samples were provided between 6 and 12 month intervals [20]. Seroprevalence estimates presented in this manuscript are based on laboratory analysis and data generated at the time of each study.

**Virus neutralization assays.** PRNTs with a semi-solid overlay were used to quantify serotype-specific neutralizing antibodies, as previously described [8]. Samples that reduced the number of plaques by 70% (i.e. PRNT70) relative to normal human serum at a serotype-specific cutoff dilution were considered positive. Cut-off dilutions were set at 1:60 for DENV-1, DENV-2, and DENV-3, and 1:40 for DENV-4 (all after the addition of virus). Positive and negative control human sera were included with each set of samples analyzed. For routine seroprevalence studies conducted in prior to 2005, positivity (e.g., 70% reduction at 1:60 dilution) was based on samples tested at 1:60 dilution (after the addition of virus). For routine seroprevalence studies conducted after 2005, positivity (e.g., 70% reduction at 1:60 dilution) was based on titers were estimated by probit regression, using a dilution series of 1:40, 1:80, 1:160, and 1:640, after the addition of virus. For quantifying end point titers, samples were diluted four-fold from 1:40 to 1:10240 and tested in duplicate; final titers were estimated by probit regression.

For seroprevalence studies, test viruses were DENV-1 16007 (DHF case from Thailand, 1964), DENV-2 16681 (Asian I genotype; DHF case from Thailand, 1964), DENV-3 IQD1728 (DF case from Peru, 2002), and DENV-4 1036 (DF case from Indonesia, 1976). DENV-2 16681 (Asian genotype) was selected for the seroprevalence assays because previous experiments in our laboratory showed that this strain minimized a cross-reactive response from DENV-1 infection [9]. For comparing antibody titers against Am-DENV-2 and AA-DENV-2 genotypes, viral strains were IQT2124 (American genotype, Iquitos, 1995), IQT2913 (American genotype, Iquitos, 1996), NFI1159 (American/Asian genotype, Iquitos 2010), and NFI1166 (American/Asian genotype, Iquitos, 2010).

Febrile surveillance. Clinic-based surveillance was conducted as described previously [17]. Briefly, participants 5 years of age or older were enrolled when reporting to outpatient clinics or hospitals in Iquitos within 7 days of onset of symptoms (fever, plus one or more other symptoms, such as headache, muscle pain, or retro-orbital pain). Within longitudinal cohorts underway during the 2010-2011 epidemic, dengue cases were identified through a community-based door-to-door active surveillance [8]. Participants’ homes were visited three times per week by technicians to inquire about residents with acute febrile illness. For both clinic-based and community-based surveillance, acute-phase serum samples were collected at time of presentation and convalescent-phase serum samples were collected two weeks to a month later. Confirmed dengue cases were defined as positive by immunofluorescence assay (IFA) in tissue culture, by reverse transcriptase polymerase chain reaction (RT-PCR), or IgM ELISA, as previously described [17]. Based on clinic-based febrile surveillance, 84% (n=433) of all DENV isolates during the 2010-2011 epidemic were DENV-2, 13% (n=69) were DENV-4 and 3% (n=13) were DENV-1.

**Contact tracing studies.** Contact tracing studies were ongoing during the 2010-2011 epidemic as part of a project aiming to capture inapparent infections and to measure the importance of human movement patterns in dengue virus transmission [21]. Upon identification of a DENV-positive febrile case through active, community-based surveillance, a cluster investigation was initiated. Participation was then solicited from all individuals living in houses recently visited by the DENV case and in neighboring households. All participants provided blood samples at days 0 and 15 to test for DENV infection by RT-PCR. Individuals participating in the cluster investigations were monitored for manifestations of febrile illness.

**Expected and observed age distributions of DENV-2 cases.** The age distribution of clinical cases is determined by the age structure of the population and at least three key epidemiological processes: immunity, the force of infection (risk of exposure) and the development of clinical symptoms. Given age-specific profiles for each of these and the age structure of the population, the expected age distribution of cases can be inferred. If we consider the scenario of a novel pathogen invading a completely susceptible population with no age dependence in risk of exposure or manifestation of disease, then the age distribution of cases would mirror the age distribution of the population. By contrast, the re-introduction of an immunizing pathogen should result in a shift of cases to lower age groups who have no prior exposure and, therefore, no protection. We would expect this shift would be less marked, however, if immunity were imperfect.

To compare the observed age distribution of DENV-2 cases in clinics and hospitals to the age distribution we computed an expected age distribution of cases based on: (1) the total number of observed cases (say N), (2) the age distribution of individuals who appeared at the clinics and hospitals with a febrile illness[17] and participated in the febrile surveillance studies in past years, and (3) age-based distribution of DENV-2 neutralizing antibodies in the months preceding the epidemic. From (2) and (3) we computed the expected probability that the next case that appears in the clinics with DENV-2 will be of a particular age (for each age group). Specifically, for each age group, we found from (2) what percent of that age group is still susceptible to DENV-2 (say, for age group *i, p_i_*) and from (3) we found what percent of those who appeared at a clinic or hospital were from that age group (say, for age group *i, a_i_*). Letting *q_i_* be the probability that the next DENV-2 case that appears at a clinic or hospital is of age group *i,* we have:

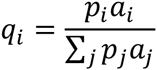

This equation is based on the assumption that an individual of any age is equally likely to become infected. By multiplying these probabilities by the total number of DENV-2 cases identified at the clinics and hospitals, we arrive at the expected number of cases by age that would have been expected if the patterns of age at time of infection were identical during the epidemic as compared with before the epidemic (specifically, for age group *i, q_i_* x *N*).

We further investigated the possibility of incomplete homologous protection against symptomatic infection by adjusting the age-based distribution of DENV-2 neutralizing antibodies (3). Above we assumed that the presence of neutralizing antibodies conferred immunity, and that only those without these antibodies could become symptomatically infected in the future. To investigate imperfect immunity, we assumed that the percentage of the individuals with neutralizing antibodies did not directly translate into the percentage of individuals who were immune. Letting *γ* be the percent of those who have protective antibodies, we recomputed *q_i_* as:

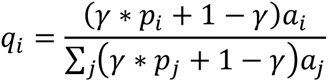

We then vary *γ* from 0 (corresponding to zero protection) to 1 (corresponding to complete protection from disease resulting from infection).

For each value of *γ*, we calculate a distribution of cases by age. We can then compare this “expected” distribution with the “observed” distribution of cases at the clinics. By calculating the probability of observing a deviation from the expected distribution as large as or larger than the one observed, we can identify which values of *γ* would not result us in concluding that there is a statistically significant difference between the expected and observed distributions. Exploiting the duality between hypothesis tests and confidence intervals, and using a cutoff of 5% for the above probability for each value of *γ*, we create a 95% confidence interval. The value of *γ* that maximizes this probability we use as our point estimate of the value of *γ* that is most supported by the data and the model. For formal comparisons, we utilize the likelihood ratio test for contingency tables (also known as the G test). For illustrative purposes, we also include analysis for the simpler Chi-square test.

## RESULTS

### The 2010-2011 DENV-2 outbreak

Based on two cohort studies that together sampled people (n=3127) in neighborhoods located throughout the city [8,20,21], in early 2010 the prevalence of neutralizing antibodies among Iquitos residents was approximately 82.1% (95% CI 80.7% -- 83.4%) for DENV-1, 74.5% (95% CI 73.0% -- 76.1%) for DENV-2, 67.1% (65.4% -- 68.7%) for DENV-3, and 40.7% (95% CI 38.9% -- 42.4%) for DENV-4 (Figure 1A), reflecting the order of DENV invasions since 1990 [17,20]. The seroprevalence of DENV-1 and DENV-2 neutralizing antibodies was strongly age-dependent (Figure 1A), as was the prevalence of multitypic neutralization profiles (neutralizing antibodies against two or more serotypes; Figure 1B). In contrast, the prevalence of DENV-3 and DENV-4 neutralizing antibodies was less age-dependent, consistent with the history of dengue outbreaks in Iquitos. The age-specific DENV-2 seroprevalence, spiking between the 10-14 and 15-19 year old age groups (Figure 1A), and analysis of cohort studies dating back to 1993 [18] strongly suggest that these titers were due to prior exposure during an Am-DENV-2 outbreak starting in 1995 rather than cross-reactive antibody for heterologous DENV serotypes.

**Figure 1.**
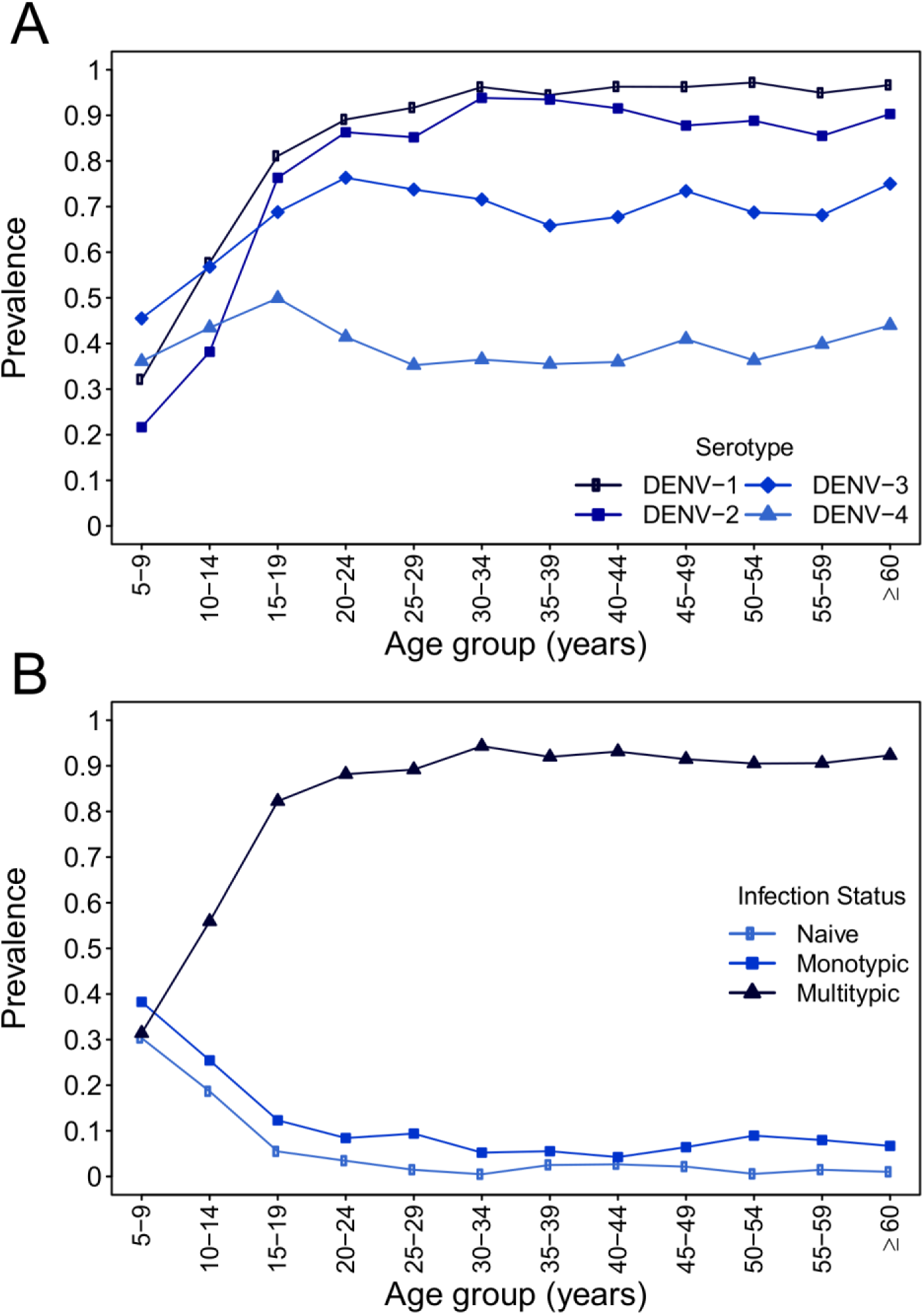
Age distribution of DENV neutralizing antibodies in 2010. Samples were collected between March and June 2010, approximately 6 months prior to a large dengue epidemic largely caused by American/Asian genotype DENV-2. Top panel: Age distribution of serotype-specific DENV neutralizing antibodies. Bottom panel: Age distribution of number of prior DENV infections. Naïve indicates absence of detectable DENV neutralizing antibodies against any serotype, monotypic indicates DENV neutralizing antibodies against one serotype, and multitypic indicates DENV neutralizing antibodies against two or more serotypes.

Given that the observed DENV-2 antibody prevalence in the Iquitos population was primarily the result of prior exposure to Am-DENV-2 ten years earlier and the sheer magnitude of the 2010-2011 dengue outbreak, we postulated that the pre-existing antibodies failed to protect against reinfection and disease. Although we did not detect any individuals with virologically-confirmed acute DENV-2 infections in both 1995 and 2010-2011, likely because of limitations of febrile surveillance activities in the mid-1990s and challenges in linking participant results across studies, we were able to test this hypothesis by analyzing population and individual level infection patterns.

### Population-level infection patterns

Age-dependent patterns of infection and disease in a population are in part a reflection of the extent of pre-existing immunity [22]. Given the high prevalence of DENV-2-neutralizing antibodies in 2010 among individuals older than 15 years (86.9%; 95% CI: 85.5% -- 88.2%; Figure 1A), we expected most new cases to occur in children. We therefore compared the observed age distribution of new AA-DENV-2 cases identified through clinic-based surveillance systems [17] with an expected distribution, based on the age-specific prevalence of DENV-2 neutralizing antibodies prior to the epidemic and the age distribution of febrile cases that participated in previous years of the studies. The observed distribution included many more cases in older age groups, especially 20 – 40 year-olds, than was expected given the age distribution of DENV-2 neutralizing antibodies prior to the introduction of AA-DENV-2 (Figure 2A). For comparison, we conducted the same analysis for DENV-4 transmission during the same time period between 2010 and 2011. For DENV-4, first introduced in 2008 and, therefore, with a much more recent history in Iquitos, the age distribution of dengue cases was consistent with what would be predicted based on age structure and detected immunity levels in the population (Figure 2B). We subsequently generated expected distributions of AA-DENV-2 infections assuming several levels of partial immunity conveyed by the pre-existing DENV-2 antibodies. We found that we could reproduce the observed distribution of DENV-2 cases when protection was incomplete (Figure 2C and 2D), with an estimated homologous protection of 35.1% (95% CI: 0% -- 65.2%) providing a best fit with the observed data.

**Figure 2.**
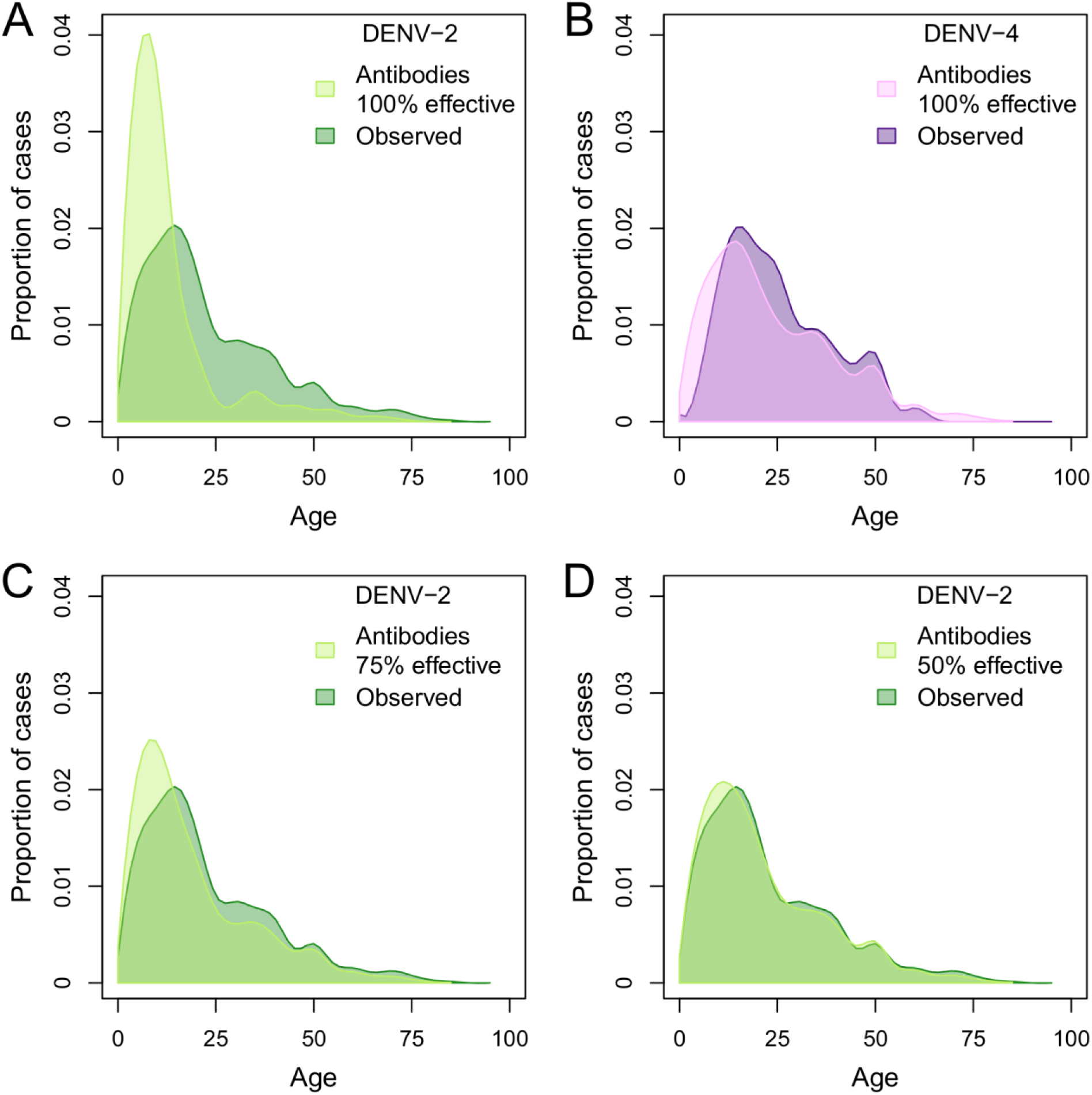
Expected versus observed DENV-2 and DENV-4 cases. The observed age distribution of cases of DENV-2 (dark green in panels A, C, and D) and DENV-4 (dark purple in panel B). Using the ages of all febrile individuals that participated in a clinic-based febrile surveillance study[17], we built an empirical estimate of the age distribution of individuals who sought treatment in Iquitos. By multiplying this distribution by the age-specific percent of the population with serotype-specific dengue antibodies, we created serotype-specific expected age distributions of cases for DENV-2 and DENV-4 (light green in panel A and light purple in panel B). We then adjusted the age- and serotype-specific immune levels and recalculated expected age distributions of cases for DENV-2 by assuming, across all ages, either 25% or 50% of those who should have been immune were still susceptible to DENV-2 (light green in panel C and light green in panel D, respectively).

The observed distribution of AA-DENV-2 cases could have been influenced by age-specific differences in infection rates. We addressed this concern using data from a case-control contact-tracing study [21] that captured inapparent DENV-2 infections and DENV-negative individuals concurrently. We compared the age-distribution of infected and uninfected individuals (Figure 3) and found them to be very similar (DENV-2 infections: mean age 31.4 years, standard deviation 19.3 years, n=75; DENV-negative: mean age 28.5 years, standard deviation 18.0 years, n=2333).

**Fig. 3.**
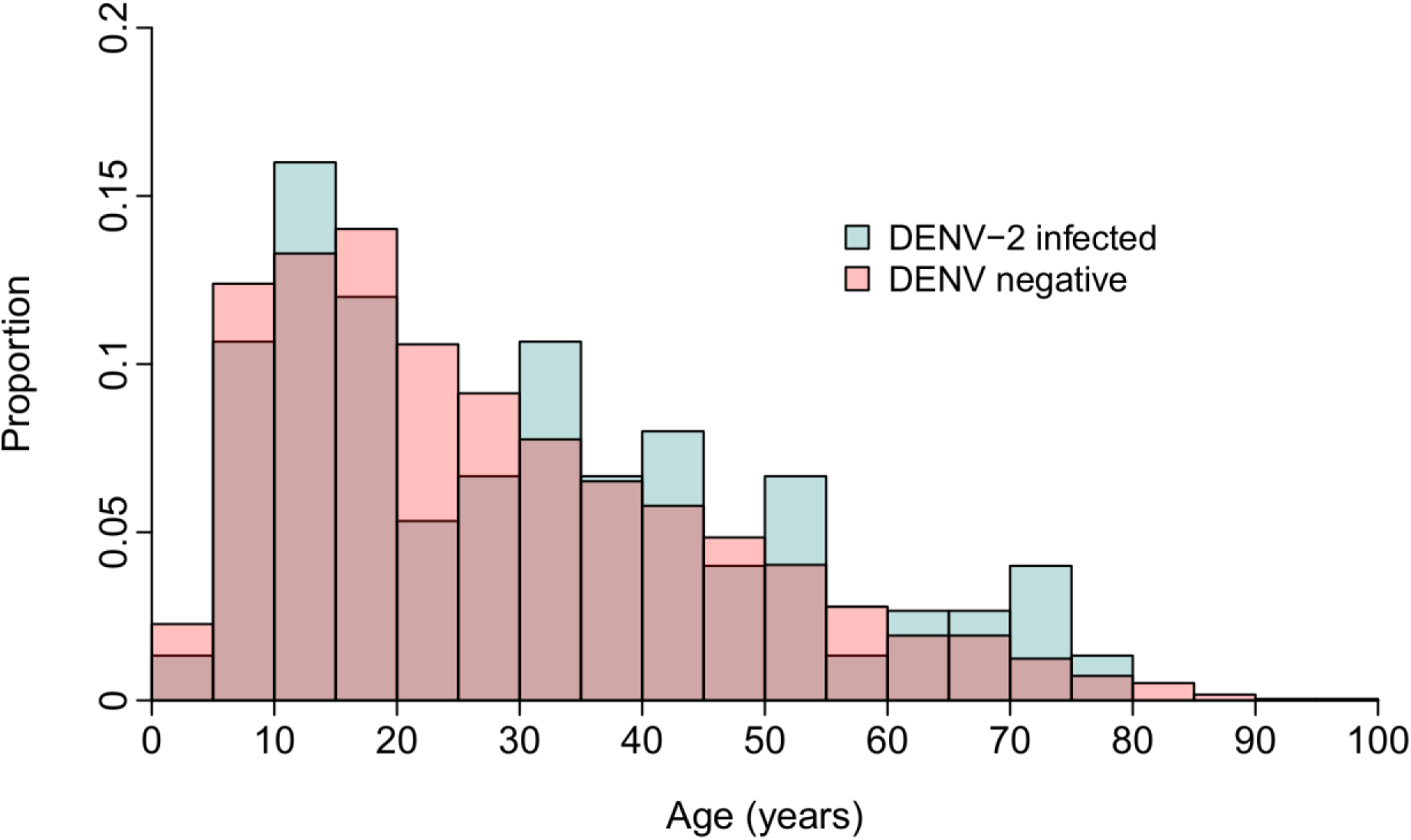
Proportion of active DENV-2-infected and DENV-negative individuals by age group (in 5 year intervals). Individuals identified through a contract-tracing study provided serum samples at day 0 and day 15 and were tested for active DENV infection by RT-PCR. Overlapping age distributions suggest that there were no gross age-dependent differences in risk for infection within the study population.

To address the possibility that the observed DENV-2 neutralizing antibodies were the result of cross-reaction from heterologous DENV infection, we utilized multiple cross-sectional samples from cohort studies conducted in Iquitos since the early 1990s [8,18,19,23]. If cross-reaction was driving the observed age dependence of DENV-2 neutralizing antibodies, we would expect that the DENV-2 antibody prevalence in adults to have increased incrementally over the years, including during periods such as 2001 – 2010 that were dominated by transmission of heterologous serotypes and near absence of confirmed clinical DENV-2 infections. Yet, among individuals born prior to 1995, the observed pattern was consistent with DENV-2 neutralizing antibodies generated largely during the 1995 outbreak (Figure 4).

**Figure 4:**
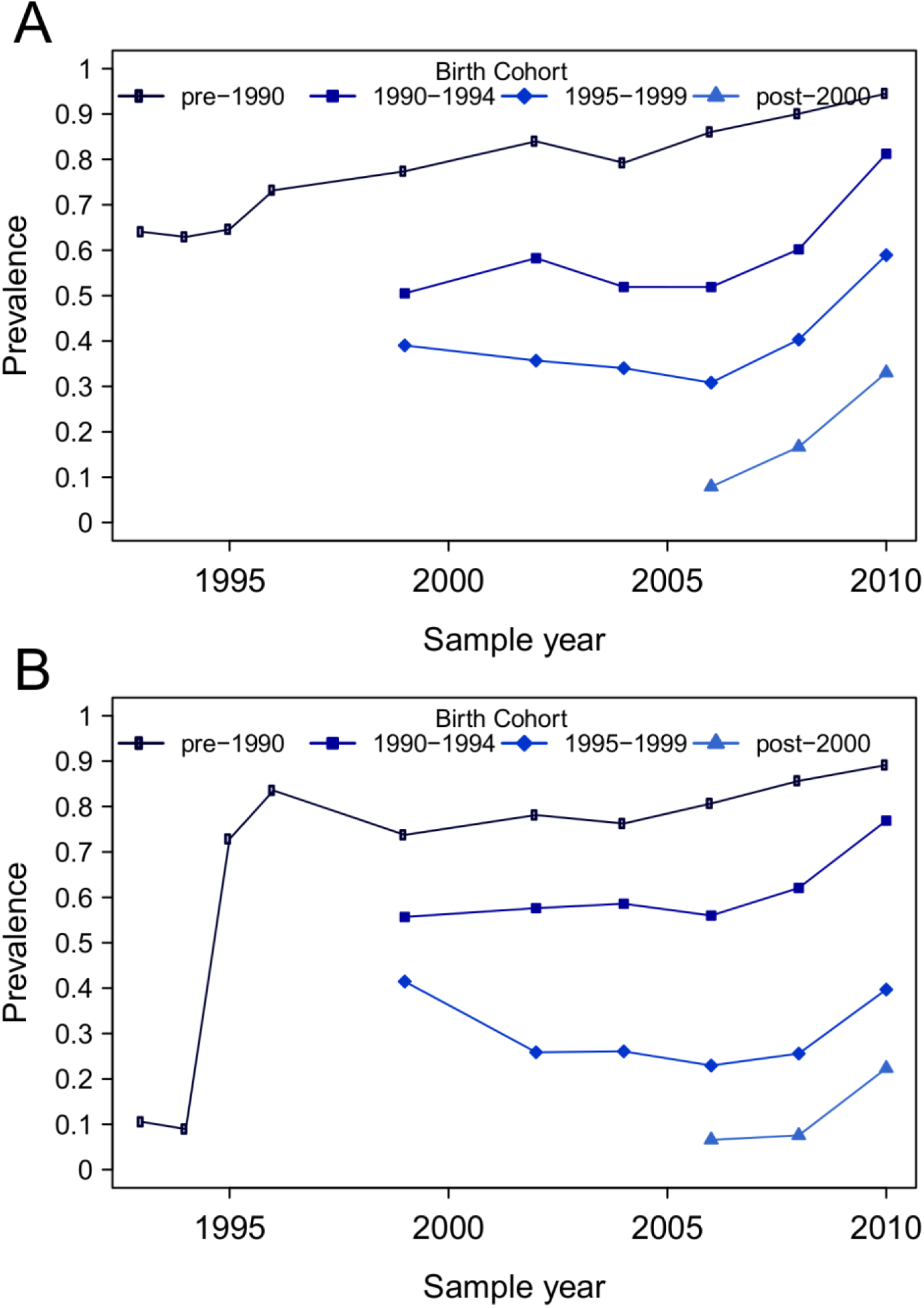
DENV-1 and DENV-2 neutralizing antibodies by birth cohort. Samples were collected from longitudinal cohort studies conducted in Iquitos since 1993. Top panel: DENV-1 neutralizing antibodies. Bottom panel: DENV-2 neutralizing antibodies.

In 1993, following the DENV-1 epidemic and prior to the known emergence of DENV-2, most of the population had DENV-1 antibodies (Figure 4A), yet fewer than 10% had DENV-2 antibodies (Figure 4B; see also [18]). Even the few DENV-2 antibodies observed prior to 1995 were likely short-term cross-reactive (heterotypic) antibodies from recent DENV-1 infection, because the majority were in DENV-1 positive individuals, and only 32% of DENV-2 antibody positive individuals maintained neutralizing antibodies during the next sampling season (i.e., from 1993 to 1994). DENV-2 antibody prevalence spiked following the 1995 DENV-2 epidemic and was largely stable among individuals born prior to 1990 (i.e., those alive during the entirety of contemporary DENV transmission in Iquitos) from 1995 to 2010 (i.e., immediately preceding the second DENV-2 epidemic; Figure 4B). For participants born after 1999 (i.e., following the first period of DENV-2 circulation), DENV-2 antibody prevalence remained low until 2010, despite experiencing more than a decade of DENV transmission (predominantly DENV-3 and DENV-4). These data indicate that DENV-2 antibodies observed in adults prior to the 2010-2011 epidemic were predominantly due to true DENV-2 infection (specifically, Am-DENV-2).

### Individual-level infection patterns

The results from the population-level analysis indicate that some individuals with pre-existing DENV-2 neutralizing antibodies prior to the 2010 epidemic were infected by AA-DENV-2 and presented with clinically apparent disease. To address this possibility, we examined the serological histories (see Methods) of individuals who had symptomatic or clinically inapparent AA-DENV-2 infections (Table 1) and who had provided two or more samples (median 4 samples; range, 2 – 7 samples) in a longitudinal cohort study. Among AA-DENV-2 cases, 43% (26/66) had consistently robust DENV-2 neutralizing titers (i.e., titers >60 for all pre-infection samples; geometric mean titer, 267) prior to the symptomatic AA-DENV-2 infection. There was a strong age dependence, consistent with the history of DENV circulation in Iquitos. Among cases born prior to 1995, 61% had DENV-2 neutralizing antibodies prior to infection, compared with 17% among those born during 1995 or later (Table 1). Among the clinically inapparent AA-DENV-2 infections, 76% (13/17) had DENV-2 neutralizing antibodies prior to infection, a prevalence similar to the study population at-large (73%) but higher than the symptomatic infections (i.e., 43%; age-adjusted odds ratio 4.2, 95% confidence interval 1.1 – 17.7). Among these AA-DENV-2-infected individuals (symptomatic and inapparent), the proportion with pre-existing DENV-2 neutralizing antibodies was consistent for years leading up to the 2010-2011 epidemic, as was the magnitude of antibody titers (Table 2). Three individuals that were monotypic for DENV-2 at the start of the study experienced virologically-confirmed acute DENV-2 infections during the 2010-2011 epidemic; two were inapparent and one was clinically apparent. These data suggest that AA-DENV-2 did infect individuals with pre-existing high titer DENV-2-neutralizing antibodies and are consistent with the notion that DENV-2 antibodies provided partial protection against disease.

**Table 1.** Pre-epidemic DENV-2 neutralizing antibody prevalence among symptomatic and inapparent DENV-2 infections.

|  | **Symptomatic** | **Inapparent** | **All** |
| --- | --- | --- | --- |
| Median age in 2010 (IQR<sup>1</sup>) | 17 (13-42) | 27 (16-45) | 19 (14-43) |
| DENV-2 antibody prevalence<sup>2</sup> | 43% (26/60) | 76% (13/17) | 51% (39/77) |
| ≤ 15yrs (born 1995 or later) | 17% (4/24) | 25% (1/4) | 18% (5/28) |
| > 15yrs (born prior to 1995) | 61% (22/36) | 92% (12/13) | 69% (34/49) |
| Geometric mean titer<sup>3</sup> | 267 | 418 | 310 |

<sup>1</sup> IQR indicates interquartile range (i.e., 25<sup>th</sup> and 75<sup>th</sup> percentiles).
<sup>2</sup> Numbers in parentheses indicate antibody positive samples and the total number analyzed in each category.
<sup>3</sup> Geometric mean titers are based on samples with detectable titers, and are aggregated across all pre-infection samples available for individuals.

**Table 2.** Pre-epidemic DENV-2 neutralizing antibody profiles of participants with subsequent confirmed DENV-2 infection in 2010–2011. Seroprevalence is based on a cutoff titer of 60. Geometric mean titers (GMT) are based on samples with detectable titers.

|  | Years prior to infection |  |  |  |
| --- | --- | --- | --- | --- |
|  | 3 to 2 yrs | 2 to 1 yrs | 1 to 0.5 yrs | ≤ 0.5 yrs |
| N<sup>1</sup> | 54 | 64 | 60 | 34 |
| Seroprevalence (95% CI) | 59% (45%-72%) | 59% (46%-72%) | 58% (45%-71%) | 59% (41%-75%) |
| Geometric mean titer<sup>2</sup> | 320 | 337 | 263 | 370 |

<sup>1</sup> If more than one sample was available for a participant within a particular time interval, only one sample was used in the calculations.
<sup>2</sup> Geometric mean titers are based on samples with detectable titers.

Our results suggest the possibility that Am-DENV-2 infection failed to generate antibodies that robustly neutralize AA-DENV-2. To address this, we utilized pre-epidemic serum collected between 2006 and 2010 from individuals who were later detected with AA-DENV-2 infections (n=21) and compared end point titers against two strains of Am-DENV-2 and two strains of AA-DENV-2 (Table 3). Robust neutralizing titers were observed against both genotypes, although pre-epidemic titers were two-fold higher against Am-DENV-2 strains (mean, 363) compared with AA-DENV-2 strains (mean, 171). In contrast, post-epidemic serum (2011-2012) collected from previously DENV-2-naïve individuals who were infected during the 2010-2011 epidemic (n=14) did not have higher titers using Am-DENV-2 test viruses (mean, 739) compared with AA-DENV-2 (mean, 899; Table 3).

**Table 3.**
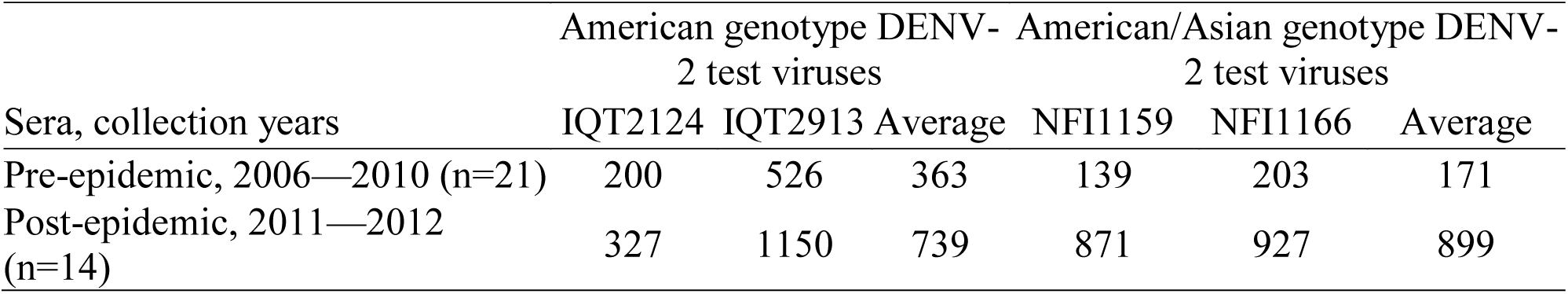
Neutralization of Am-DENV-2 and AA-DENV-2 test viruses using human serum from longitudinal cohort studies. Geometric mean titers (GMTs) for the individual test viruses and the average of the two GMTs are shown.

## Discussion

We provide indirect evidence that challenges a fundamental assumption of dengue immunology: that infection with one serotype conveys lifelong protection to reinfection by all strains belonging to the same serotype. A newly introduced genotype of DENV-2 exhibited characteristics similar to a novel serotype in a population with a high prevalence of DENV-2 neutralizing antibodies. Moreover, we isolated DENV-2 virus from numerous individuals with pre-existing, robust DENV-2 antibody titers. Epidemiological data from long-term longitudinal cohort studies[20] show that pre-existing DENV-2 antibodies present in the adult population were mostly a product of the invasion of an American strain of DENV-2 in 1995. Although these antibodies did not appear to completely protect against infection with the new strain, they did appear to provide partial protection against febrile illness, reducing the probability of disease. Dengue epidemiology and vaccine development have long been complicated by the immunological interplay among serotypes at the level of the human host. Genetic and phenotypic heterogeneity may be sufficient to extend this interplay to the level of individual genotypes, which markedly increases the complexity of the system[7] and could present further challenges for the development of a suitable dengue vaccine[12,24].

One key piece of evidence in our study arises from the observation of more infections than expected in older age groups. Our calculations for the expected age distribution of DENV-2 cases take into account the age-dependent participation in our febrile illness studies. Our previous analysis of DENV-3 and DENV-4 seroconversions did not indicate an substantial age-dependent risk for infection [8]. Further, among confirmed infections, we did not observe a marked age-dependent difference in risk for disease for DENV-3, and we observed only a modest increase in disease risk among younger adults for DENV-4 [8]. Although others have reported that disease risk may increase with age [25], this conclusion is based on indirectly measured exposures and could be skewed by reporting biases. Additionally, in Iquitos, the large majority of individuals older than 20 years (~90%; Figure 1B) have been exposed to at least two serotypes, which would be expected to dramatically reduce the risk of disease upon infection relative to younger participants [8]. Therefore, we feel that our assumptions are conservative and likely underestimate the relative number of cases among older participants.

Our results depend in part on PRNTs, which are widely considered the most specific serological tests available for DENV. Yet even PRNTs can be prone to cross-reaction in some circumstances [26]. To reduce potential for cross-reaction, we used more stringent conditions (PRNT70 at 1:60 dilution cutoff) than used in most dengue vaccine [12] or dengue epidemiological studies [27], which often use a 50% reduction (PRNT50) threshold. We conducted an additional evaluation of even more stringent conditions (90% reduction threshold, or PRNT90) had only modest reductions in seroprevalence (67% for PRNT90 at a 1:60 dilution cutoff versus 73% for PRNT70 as reported above). Furthermore, among individuals who had pre-existing DENV-2 neutralizing antibodies and were infected during the 2010-2011 epidemic, more than half had titers greater than 300 (using 70% reduction threshold). This titer (323, using the lower 50% reduction threshold) was proposed as a cutoff for differentiating “susceptible” from “non-susceptible” subjects based on limited data from a Thai cohort [28].

More importantly, we directly address the issue of cross-reaction by utilizing samples collected prior to the first documented DENV-2 outbreak in 1995. Detectable DENV-2 antibody prevalence was low prior to DENV-2 emergence in Iquitos, spiked immediately following DENV-2 emergence in 1995, and remained stable among adults up to the 2010-2011 epidemic (Figure 4). Further, the prevalence of antibodies in individuals born after the major DENV-2 transmission period (1995—1999) was low, despite potential for cross-reaction following epidemics of DENV-3 and DENV-4 between 2001 and 2009. The absence of a large age shift in overt disease during the AA-DENV-2 epidemic also supports our conclusion. In contrast, the 2007-2008 re-emergence of DENV-2 in Brazil was accompanied by an age shift in dengue cases, with a sharp increase in disease incidence in younger age groups relative to older individuals [29]. Notably in the case of Brazil, AA-DENV-2 had circulated during earlier outbreaks (before 2002) prior to re-emerging in 2007.

It is not clear if the homologous serotype re-infection we identified is unique to the DENV-2 genotypes we studied (i.e., American genotype followed by American/Asian genotype) or if it is a more widespread phenomenon among other dengue viruses. Within the E gene coding region, there is approximately 10% nucleotide divergence and 2% amino acid divergence between Am-DENV-2 and AA-DENV-2. Am-DENV-2 has been shown to differ from Asian and American/Asian genotypes phenotypically, including replication in tissue culture and incubation periods in mosquitoes[30,31]. Further, for other arthropod-borne viruses and influenza viruses, a few amino acid changes can have profound antigenic and epidemiological consequences [32]. There is some support for re-infection with a homologous DENV serotype in animal and in vitro studies, although most studies do not utilize naturally occurring virus variants and do not challenge hosts with a different genotype of a homologous serotype. In vaccine studies in non-human primates, viremia may be attenuated but detectable despite the presence of a robust immune response [33]. Even when viremia is undetectable, sufficient viral replication occurs to stimulate homotypic antibody boosting [10]. In one challenge study, four of five monkeys had detectable viremia upon challenge with an Asian genotype of DENV-2 following primary infection by different DENV-2 genotypes [34]. For virus neutralization studies using sera from vaccinated and naturally infected humans [9,27], monkeys, and mice, there are often marked genotype-dependent differences in neutralization capacity [11], some greater than 100-fold. Similarly, monoclonal antibodies vary greatly in their neutralizing activity in cell culture and protective efficacy in mice [11]. We found quantitative, but not qualitative, differences in neutralization of Am-DENV-2 compared with AA-DENV-2 (Table 3). Together with our data, these neutralization results support the need to re-evaluate underlying assumptions about the epidemiological importance of genotypic differences in neutralization capacity [11].

To our knowledge, this is the first study to present epidemiological data as evidence for homologous reinfection by DENV. Numerous other studies have noted a poor correlation between pre-existing virus neutralization antibody titers and clinical protection, both in natural infection [27,28] and vaccine trials [12,35]. For example, in a Thai cohort [27], pre-existing DENV-2 antibodies were detected in 60% of DENV-2 infections. Notably, there was one DENV-2 case with pre-existing monotypic PRNT pattern for DENV-2, suggestive of re-infection. In an ongoing clinic-based surveillance system in Puerto Rico, one individual had two distinct DENV-2 episodes, detected by RT-PCR, 16 months apart [36]. It is possible that homologous reinfection has occurred in other settings and our awareness of this phenomenon is limited by study design and diagnostics constrains. For example, genotype replacement events are not commonly documented in detail [7]. Moreover, unlike our studies in Iquitos, most community-based cohort studies are short term or monitor only children [7,27], thus it is not possible to carryout long-term follow-up across different age groups. Most available long-term dengue datasets include only severe, hospitalized cases, which may not be representative of actual virus transmission dynamics. However, an analysis of 323 virologically confirmed sequential infections in Thailand, based largely on 17 years of hospital-based passive surveillance, did not detect cases of homologous re-infection [37]. Further, homologous challenge studies in humans conducted in the 1940s by Albert Sabin did not find evidence for re-infection. The Sabin challenge studies were conducted, however, over a short time frame (< 18 months) with a small sample of participants and utilized the same DENV-2 strain for initial infection and challenge [3]. In our study we did not detect any individuals with virologically-confirmed acute DENV-2 infections in both 1995 and 2010-2011, likely in large part because of limitations of febrile surveillance activities in the mid-1990s and challenges in linking participant samples from across studies and years. Subsequent studies should be designed to track individuals’ DENV infections over longer time periods to confirm our findings and to determine the relevance of re-infection to DENV transmission dynamics.

### Conclusions

Consistent with other reports [24,27,28], our data demonstrate that the presence of detectable neutralizing antibodies does not necessarily correlate with protection from a homologous serotype challenge Although cross-neutralizing antibodies do arise from heterologous serotype infection [11], our data indicate that the majority of neutralizing antibodies were from prior infection and thus our results are consistent with widespread homologous re-infection by DENV-2. Our results have direct implications for current approaches for design and development of dengue vaccines. In recent phase 2b and phase 3 trials, a high proportion of vaccinated individuals developed neutralizing antibodies, with geometric mean titers exceeding 140 for all serotypes, yet protective efficacy against DENV-2 was far from complete (<50%) [12,24]. Although viral interference has been suggested as one explanation for the vaccine failure, our results indicate that success of a vaccine based on a single strain of each serotype may be dependent on circulating strains. There is an urgent need for better *in vitro* measures of protection, because positive PRNT results do not accurately predict protection from symptomatic infection, particularly at the liberal cutoffs used in vaccine immunogenicity studies (e.g., 50% reduction with 1:10 dilution of serum).

## Acknowledgements

We thank the study participants in Iquitos for their time and willingness to participate. We thank the numerous field and laboratory personnel in Lima and Iquitos, particularly Carolina Guevara, Zonia Rios, Wieslawa Alava, Rebecca Carrion, Leslye Angulo, and Guadalupe Flores. Authors Eric S. Halsey and Tadeusz Kochel are military service members and Angelica Espinoza and Stalin Vilcarromero are employees of the U.S. Government. This work was prepared as part of their official duties. Title 17 U.S.C. § 105 provides that ‘Copyright protection under this title is not available for any work of the United States Government’. Title 17 U.S.C. § 101 defines a U.S. Government work as a work prepared by a military service members or employees of the U.S. Government as part of those persons’ official duties. The views expressed in this article are those of the authors and do not necessarily reflect the official policy or position of the Ministries of Health of Peru or Department of the Navy, Department of Defense, nor the U.S. Government.

